# Slow recovery from inbreeding depression generated by the complex genetic architecture of segregating deleterious mutations

**DOI:** 10.1101/862631

**Authors:** Paula E. Adams, Anna L. Crist, Ellen M. Young, John H. Willis, Patrick C. Phillips, Janna L. Fierst

## Abstract

The deleterious effects of inbreeding have been of extreme importance to evolutionary biology, but it has been difficult to characterize the complex interactions between genetic constraints and selection that lead to fitness loss and recovery after inbreeding. Viruses, bacteria, and the selfing nematode *Caenorhabditis elegans* have been shown to be capable of rapid recovery from the fixation of novel deleterious mutation, however the potential for fitness recovery from fixation of segregating variation under inbreeding in outcrossing organisms is poorly understood. *C. remanei* is an outcrossing relative of *C. elegans* with high polymorphic variation and extreme inbreeding depression. Here we sought to characterize changes *C. remanei* in patterns of genomic diversity after ∼30 generations of inbreeding via brother-sister mating followed by several hundred generations of recovery at large population size. As expected, inbreeding led to a large decline in reproductive fitness, but unlike results from mutation accumulation experiments, recovery from inbreeding at large populations sizes generated only very moderate recovery in fitness after 300 generations. At the genomic level, we found that while 66% of ancestral segregating SNPs were fixed in the inbred population, this was far fewer than expected under neutral processes. Under recovery, 36 SNPs across 30 genes involved in alimentary, muscular, nervous and reproductive systems changed reproducibly across all replicates, indicating that strong selection for fitness recovery does exist but is likely mutationally limited due to the large number of potential targets. Our results indicate that recovery from inbreeding depression via new compensatory mutations is likely to be constrained by the large number of segregating deleterious variants present in natural populations, limiting the capacity for rapid evolutionary rescue of small populations.

**Impact Summary:** Inbreeding is defined as mating between close relatives and can have a large effect on the genetic diversity and fitness of populations. This has been recognized for over 100 years of study in evolutionary biology, but the specific genomic changes that accompany inbreeding and the loss of fitness are still not known. Evolutionary theory predicts that inbred populations lose fitness through the fixation of many deleterious alleles and it is not known if populations can recover fitness after prolonged periods of inbreeding and deleterious fixations, or how long recovery may take. These questions are particularly important for wild populations experiencing declines. In this study we use laboratory populations of the nematode worm *Caenorhabditis remanei* to analyze the loss of fitness and genomic changes that accompany inbreeding via brother-sister mating, and to track the populations as they recover from inbreeding at large population size over 300 generations. We find that:

1. Total progeny decreased by 65% after inbreeding
2. There were many nucleotides in the genome that remained heterozygous after inbreeding
3. There was an excess of inbreeding-resistant nucleotides on the X chromosome
4. The number of progeny remained low after 300 generations of recovery from inbreeding
5. 30 genes changed significant in allele frequency during recovery, including genes involved in the alimentary, muscular, nervous and reproductive systems

Together, our results demonstrate that recovery from inbreeding is difficult, likely due to the fixation of numerous deleterious alleles throughout the genome.

## Introduction

“The evil effects of close interbreeding” have been of importance to geneticists and evolutionary biologists since Darwin first wrote about them in 1896 (Darwin 1896). Inbreeding depression is defined as the reduction in fitness incurred from reproduction between closely related individuals (Charlesworth and Charlesworth 1987). This reduced fitness can lead to decreased fecundity and eventual extinction of small populations (Hedrick and Garcia-Dorado 2016). Inbreeding can have a large effect on the success of conservation of endangered or isolated species (Kardos et al. 2016). However, despite a developed understanding of the significance of inbreeding depression, identifying specific alleles contributing to the reduction in fitness has remained a challenge (Hedrick and Garcia-Dorado 2016). From a conservation point of view, we know even less about the likelihood that populations that have undergone a history of inbreeding can recover in fitness via contributions of new adaptive mutations (Hedrick and Kalinowski 2000). In this sense, inbreeding shifts the population from its current fitness optimum and new mutations or other forms of genetic input are needed to “rescue” the population from continued degradation in fitness (Whitlock and Otto 1999; Whitlock et al. 2003; Gonzalez et al. 2013; Bell et al. 2019). What is the genetic basis of inbreeding depression and is it possible for a population to recover from the deleterious effects of inbreeding after it has occurred? Here, we address these questions by characterizing fitness reduction and genomic changes in the nematode worm *Caenorhabditis remanei* after inbreeding and throughout recovery at large population sizes.

During inbreeding, large regions of the genome can become homozygous. Inbreeding depression can be caused by an accumulation of recessive deleterious alleles that fix during inbreeding or by the fixation of segregating alleles at loci in which heterozygotes have a fitness advantage (Charlesworth and Charlesworth 1987; Charlesworth and Willis 2009). Mutation accumulation studies have attempted to characterize the spectrum of deleterious alleles (Charlesworth et al. 1993). Theory suggests that most mutations are slightly deleterious, and over time genetic drift in small or inbred populations will lead to fitness declines as slightly deleterious alleles accumulate (Lande 1994; Lynch et al. 1999; Lynch and Gabriel 1990). For example, in the self-reproducing *C. elegans* this decline is on the order of 0.1% per generation (Vassilieva et al. 2000) while outcrossing *Caenorhabditis* experience more rapid fitness decay (Baer et al. 2010). Interestingly, populations that have experienced recent fixation of novel deleterious mutations are able, for the most part, to rapidly recover and return to their initial fitness state within a few dozen generations (Estes and Lynch 2003; Estes et al. 2004; Estes et al. 2011), likely due to compensatory mutations at other sites in the genome (Denver et al. 2010). Similar observations have been made in other systems (Burch and Chao 1999; Whitlock and Otto 1999; Maisnier-Patin et al. 2002). These observations raise the possibility that genetic rescue of inbred populations via compensatory mutation might not particularly difficult, as the total number of potential compensatory sites is in principle very large.

However, inbreeding depression in most populations is likely generated by the accumulation of segregating deleterious mutations over a long period of time and potentially at a large number of loci. Thus, while the effects observed in mutation accumulation studies are the ultimate source of inbreeding depression in natural populations, they may not reflect the long-term segregating effects of mutations that have been filtered through population-level processes of natural selection, genetic drift and genomic linkage. Indeed, inbreeding assays of natural isolates have shown minimal fitness loss in the self-reproducing *C. elegans* but very severe fitness loss and line-specific extinction up to ∼90% in the outcrossing *C*. *remanei* (Dolgin et al. 2007), with the difference almost certainly driven by the likelihood that deleterious recessive mutations will be exposed to natural selection under these two mating systems (Lande and Schemske 1985). Thus, while we expect that inbred populations *can* recover after the fixation of deleterious mutations (Estes and Lynch 2003; Denver et al. 2010; Estes et al. 2011), whether they *will* recover following the fixation of segregating variants is an open question.

Historically, pedigree information has been used to predict the probability of a diploid allele being identical-by-descent (IBD) (Hedrick and Garcia-Dorado 2016). Large IBD runs of homozygosity (ROH) can be detected in sequence data and then used to infer the amount of inbreeding in the absence of pedigree information (Kardos et al. 2016; Hedrick and Garcia-Dorado 2016). Larger IBD segments indicate more recently related ancestors, whereas short IBD segments indicate more distantly related common ancestors on average (Kardos et al. 2016). Using these methods, whole-genome sequencing can be used to characterize the amount of inbreeding within a population and to identify regions of any potential genetic resistance to inbreeding (e.g., because of overdominance). However, identifying the specific alleles underlying fitness loss and genetic resistance has remained a challenge (Hedrick and Garcia-Dorado 2016).

Here, we use whole-genome sequencing in *C. remanei* to first study allelic changes that accompany fitness loss through inbreeding and to second track genetic changes in replicate populations over 200 generations as they recover from this inbreeding in very large populations. Analyzing the first phase of inbreeding allows us to quantify how many loci were fixed during this process, as well as how many displayed resistance to inbreeding. Analyzing the second phase of recovery from inbreeding allows us to observe genomic changes that are parallel across recovery lines. Our results show that, in contrast to expectations generated from mutation accumulation experiments, fitness recovery from inbreeding may not be so easily accomplished because of the scope and scale of segregating deleterious genetic variation within natural populations.

## Methods

### Inbreeding

To overcome the extinction reported for *C. remanei* (Dolgin et al. 2007) a novel scheme was used for inbreeding (hereafter referred to as “Inbred”; Fierst et al. 2015). *C. remanei* strain EM464 (hereafter referred to as “Ancestor”) was originally isolated in New York City and obtained from the Caenorhabditis Genetics Center, University of Minnesota, Minneapolis, MN. Two hundred independent lines of the Ancestor were subjected to brother-sister mating with just 2 lines remaining at generation 7. These lines were maintained for 20 generations as an outcrossing population. From this population 100 lines were subjected to brother-sister mating for 23 generations until only one surviving Inbred line, PX356, remained (Fig. 1; (Fierst et al. 2015).

**Figure 1.**
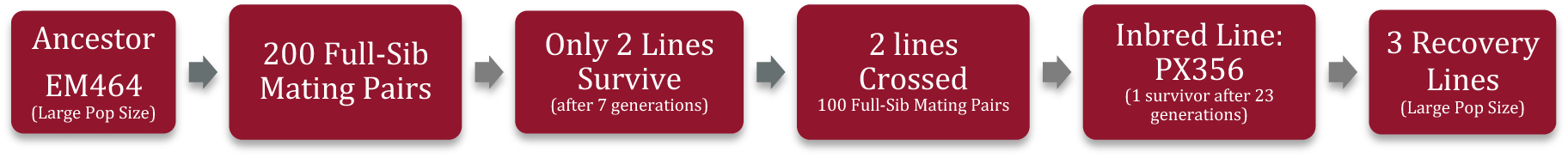
The inbreeding and recovery scheme used to create the Inbred line from the Ancestral strain of *C. remanei*. Two hundred plates with full-sibling mating pairs were kept through 7 generations until only 2 remained alive. Those 2 lines were allowed to expand for 20 generations then crossed to create 100 full-sib mating pairs. These lines were transferred for 23 generations until only 1 line, the Inbred PX356, was left alive. Offspring of the Inbred line were allowed to reproduce at large population size in 3 replicate Recovery lines for 300 generations.

### Maintenance of Recovery Lines

Three Recovery lines were independently established from the Inbred line (details of laboratory culture and experimental set-up are given in the Supplementary Methods). Recovery lines were propagated by transferring a piece of agar from a populated petri dish and placing it upside down on the agar surface of a new petri dish every 3-4 days. Each transfer event was counted as one generation and populations grew to census sizes of >2,000 individuals in-between transfers.

### Experimental Assays for Fecundity and Longevity

After inbreeding and recovery, fecundity and longevity assays were conducted on population samples. The Inbred line was included in each experiment as a control. To measure fecundity, 40 replicates of each line containing 1 virgin L4 female and 3 L4 males were established. Every 24 hours for 1 week, the worms were transferred to new 35 mm agar plates. The plates the worms were transferred from were kept for 2 days, after which L4 progeny were counted and deaths recorded.

To measure longevity, 30 replicates containing 5 virgin L4 females were established. Plates were examined every 1-2 days to check for dead individuals. Individuals were transferred to new petri dishes on day 10 of the experiment and every 7 days after that to ensure adequate amounts of the bacterial food source and to avoid contamination.

### DNA Isolation

DNA was isolated from pooled population samples and sequenced on an Illumina HiSeq instrument. Recovery Lines 1, 2 and 3 were sequenced as single end DNA reads after 100 generations and Recovery Line 2 was sequenced as single end DNA reads after 200 generations. Recovery Lines 1 and 3 were sequenced as paired end DNA reads after 200 generations.

### Genetic Analyses

DNA libraries were aligned to the PX356 reference sequence NMWX00000000.1 using 2 alignment softwares, GMAP-GSNAP (Wu et al. 2016) and BWA mem (Li and Durbin 2009). Picard Tools (Institute 2016) and the Genome Analysis Toolkit (GATK) were used to filter noise in alignment (DePristo et al. 2011; McKenna et al. 2010) and the software package MAPGD used to estimate allele frequencies and identify segregating variants (Lynch et al. 2014; Ackerman et al. in prep). Alignments were filtered for coverage (all bioinformatics scripts and workflows are available at https://github.com/BamaComputationalBiology/Inbreeding). The minimum sequence read coverage was 5 for the Ancestor and Recovery lines and 10% of the mean coverage (37 sequence reads) for the Inbred line. The maximum coverage was 3x the mean coverage for all lines (Supplementary Table 1(Li 2014). RepeatMasker was used to identify repeat regions (Smit et al. 2013-2015) and repeat-associated SNPs excluded from analyses.

The Inbred line was sequenced at a high mean read depth of 370x while the Ancestor and Recovery lines were sequenced to mean depths of 25-64x (S. Table 1). After filtering, 150,348 sites (0.13% of the 118.5Mb assembled genome) displayed segregating variants.

### Allele Frequency Estimation

Allele frequencies were estimated with the MAPGD software package (Ackerman et al. in prep; Lynch et al. 2014). Sites with missing data were removed and SNPs with a log-likelihood ratio >22 and a minor allele frequency >5% were considered to be true segregating variants. We required segregating sites to meet these criteria for both BWA (Li and Durbin 2009) and GSNAP (Wu et al. 2016) alignments to reduce false positives and remove sites with ambiguous alignment (Kofler et al. 2016) and used the BWA allele frequencies in analyses.

Because our data were a somewhat heterogeneous combination of paired end and single end sequences at different read depths, we sought to remove potential biases. In particular, segregating polymorphisms were increased in both paired end and high depth samples (S. Table 1) and we removed nucleotides with segregating variants in paired-end sequences that displayed fixation (no polymorphism) in the single-end samples. These sites may have been true polymorphisms, but with our design they could not be distinguished from sampling error. We calculated the Site Frequency Spectrum (SFS) for each sample using the minor allele frequencies at each variable site (Fisher 1930; Wright 1938).

### Runs of Homozygosity

We defined a run of homozygosity (ROH) as a region of the genome greater than 1kb in length where minor allele frequency did not exceed 3%. This was roughly the threshold of detection (equivalent to 1-2 sequence reads) for our samples that were sequenced as single end reads. This procedure eliminates small ROH and may underestimate the size of ROH and we chose to take this approach to focus on genome-wide patterns for which we had rigorous support.

### Allele Frequency Trajectories

We separated nucleotides by allele frequency trajectories to identify the major trends occurring during inbreeding and recovery. ‘Fixation’ nucleotides were defined as segregating in the Ancestor and >95% major allele frequency in all Inbred and Recovery lines. ‘Intermediate’ were those segregating in the Ancestor and changed in frequency <50% through inbreeding and recovery. The remaining sites were filtered into four trends: (1) ‘Bounce Down’ sites had low frequency in the Ancestor, higher frequency in the Inbred, and lower frequency in recovery; (2) ‘Up’ sites increased in frequency during both inbreeding and recovery; (3) ‘Bounce Up’ sites had high initial frequency, lower frequency during inbreeding and higher frequency during recovery; and (4) ‘Down’ sites had high frequency in the Ancestor that decreased through inbreeding and recovery. These categories allow us to characterize what proportion of variable nucleotides were fixed through inbreeding and, of the remaining nucleotides, how segregating variation changed through recovery.

### Effective Population Size

Effective population sizes were calculated with the software package PoolSeq (Taus et al. 2017). Census size was varied from 1500 (the approximate population size of a plate of nematodes) to 1,000,000 (the estimated effective population size for the species (Cutter et al. 2006) to test the influence of parameters on effective population size estimation.

### Selection scans in recovery lines

We used two methods to identify significant allele frequency changes in Recovery lines. First, we fit a general linear model (GLM) with quasibinomial error distribution to the allele frequency changes across the Inbred line, generation 100 Recovery, and generation 200 Recovery according to the Wiberg et al. (2017) recommendation for best practices with pooled sequencing data. Second, we performed a Cochran-Mantel-Haenszel (CMH) test to analyze parallel changes in allele frequencies between the Inbred and Recovery lines at generation 100 and 200 with the software package PoPoolation2 (Kofler et al. 2011). All sites that were significant in the quasibinomial-GLM analyses were also significant with the CMH test and we retained all significant sites for analysis. We used the R software package *qvalue* for false discovery rate correction (Storey et al. 2019). Nucleotides with significant changes (i.e., quasibinomial-GLM qvalue < 0.05) across all three Recovery lines were associated with genic or intergenic locations with BEDTools (Quinlan and Hall 2010). Proteins containing significant SNPs were annotated for putative molecular functions with the Interproscan software package (Jones et al. 2014) and orthologous genes in other *Caenorhabditis* species identified with OrthoFinder (Emms and Kelly 2015). We searched WormBase ParaSite for functional information for orthologous genes (Howe et al. 2017).

### F_ST_

We used the software package PoolFstat (Hivert et al. 2018) to calculate the fixation index (F_ST_) between population pairs for each variable SNP. We calculated the mean F_ST_ for each gene by averaging across variant sites 1kb upstream of the gene, within the gene and 1kb downstream of the gene.

## Results

### Fecundity and Longevity

The mean cumulative per individual progeny for the Ancestor was 563 ± 35 and inbreeding decreased this to 196 ± 8, a 65% reduction (Fig. 2A). Total progeny increased by 44% to 283 ± 12 after 200 generations of recovery but unexpectedly shrank to 219 ± 12 after another 100 generations of recovery (Fig. 2A). In contrast, the mean lifespan in the Recovery lines was 4 days longer than that of the Ancestor and the oldest individual in the Recovery lines lived 12 days longer than the longest living Ancestor (Fig. 2B).

**Figure 2.**
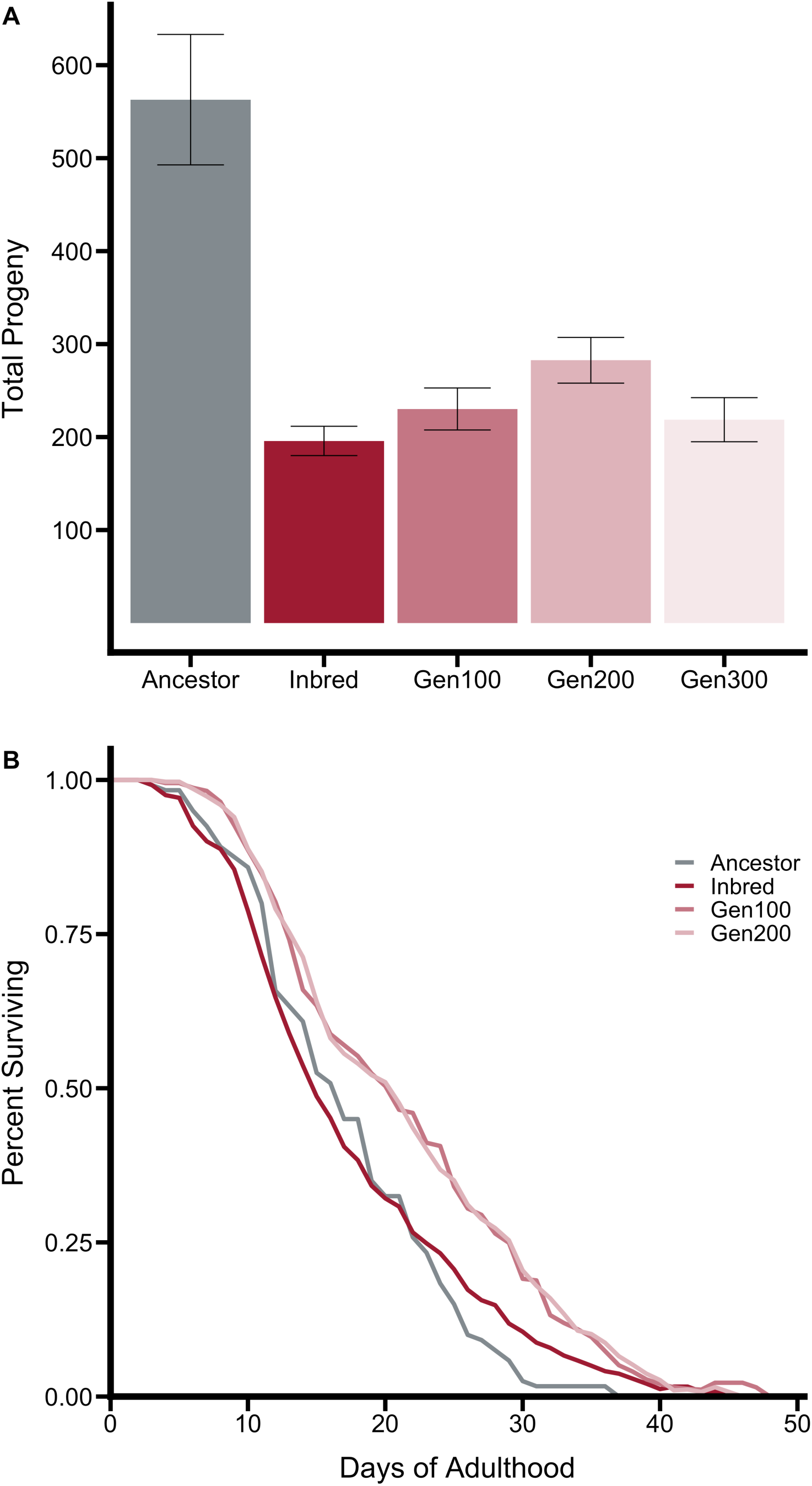
The phenotypic effects of inbreeding included (A) a decrease in the mean reproductive output that was not recovered after 300 generations of breeding at large population sizes. There was (B) no influence of inbreeding on longevity but the Recovery lines evolved an increase in longevity when compared with the Ancestral and Inbred lines.

Age-specific fecundity differed among lines (Fig 3). The Inbred line completed 90% of its egg laying within the first 3 days of reproduction and 100% of its egg laying within 5 days. In comparison, the Ancestor completed 52% of its egg laying within the first 3 days of reproduction and continued egg laying at a low rate for the 7 day assay period. The Recovery lines completed 76-81% of their egg laying within the first 3 days and continued egg laying at decreasing rates for 7 days.

**Figure 3.**
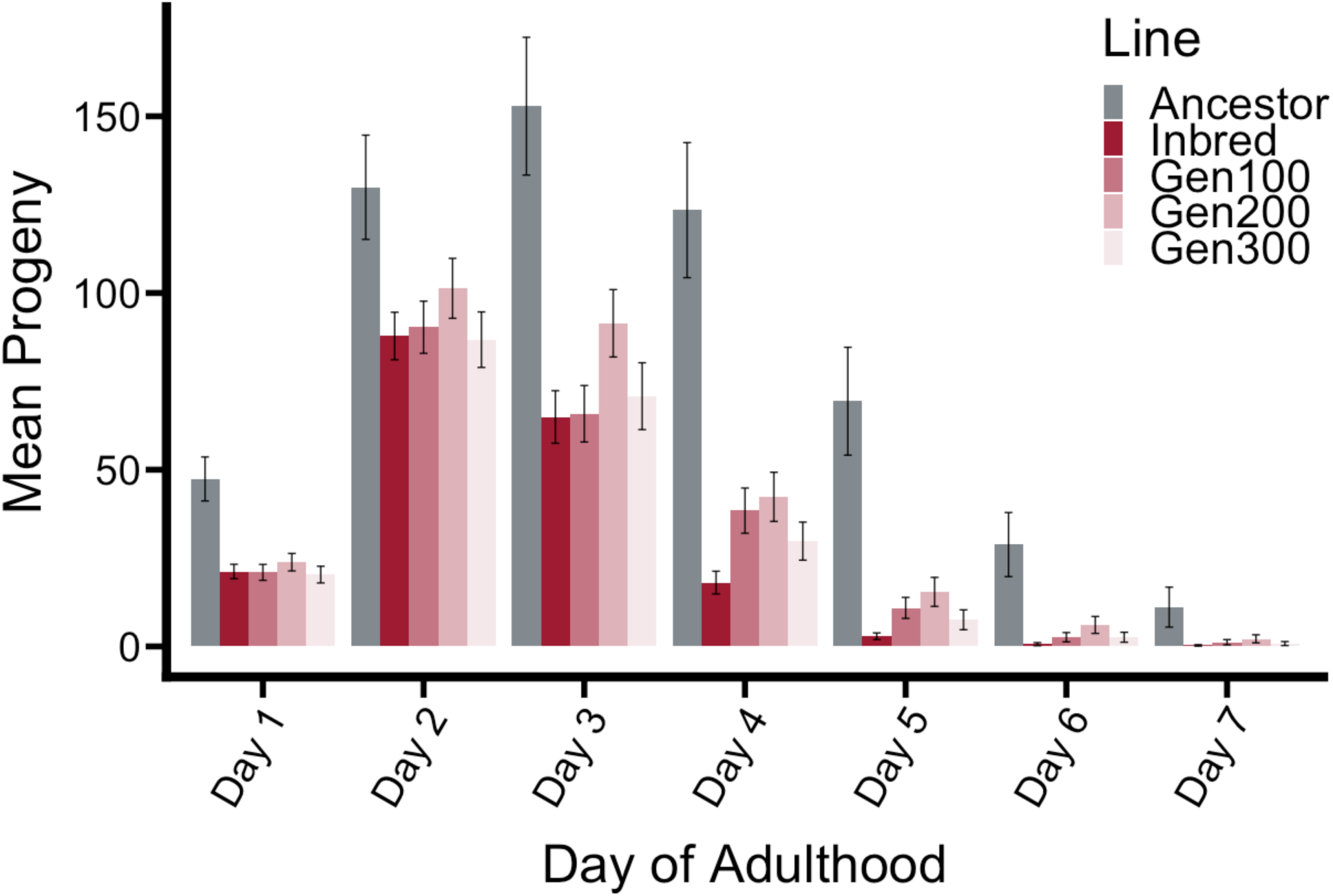
Mean progeny by day of adulthood.

### Allelic Diversity

Allelic diversity was reduced during inbreeding (Fig. 4A-E). Of the 150,348 segregating sites observed, 139,658 (93%) of these were variable in the Ancestor and 51,408 (34%) were variable in the Inbred line. It is difficult to exactly calculate a neutral expectation for homozygosity under our inbreeding design because the brother-sister mating was paused at generation 7 and then continued for an additional 23 generations (Fig. 1). However, we can use 23 generations of inbreeding as a minimum for our homozygosity expectation, noting that the true expectation will be somewhere between 23 and 30 generations of inbreeding. On average we expect brother-sister mating to homogenize ½ of the heterozygous variants each generation and (½)^23^ or ∼1.2×10^-7^ will remain after inbreeding. With a starting point of 139,658 segregating sites in the Ancestor we would expect, on average, 0.017 SNPs to remain after this period of inbreeding. Our actual number of segregating variants in the Inbred line, 51,408, is far from this neutral expectation and indicates multiple inbreeding-resistant sites (Barrière et al. 2009). The Recovery lines had an average of 45,853 sites with segregating variants (30% of the total) in generation 100 and 50,593 segregating sites (34% of the total) in generation 200. This is likely an underestimate of true segregating diversity in recovery due to the high sequence depth of our Ancestor and comparatively low sequence depth of our Recovery lines, but it demonstrates that little genetic variation was regained or generated in recovery.

**Figure 4.**
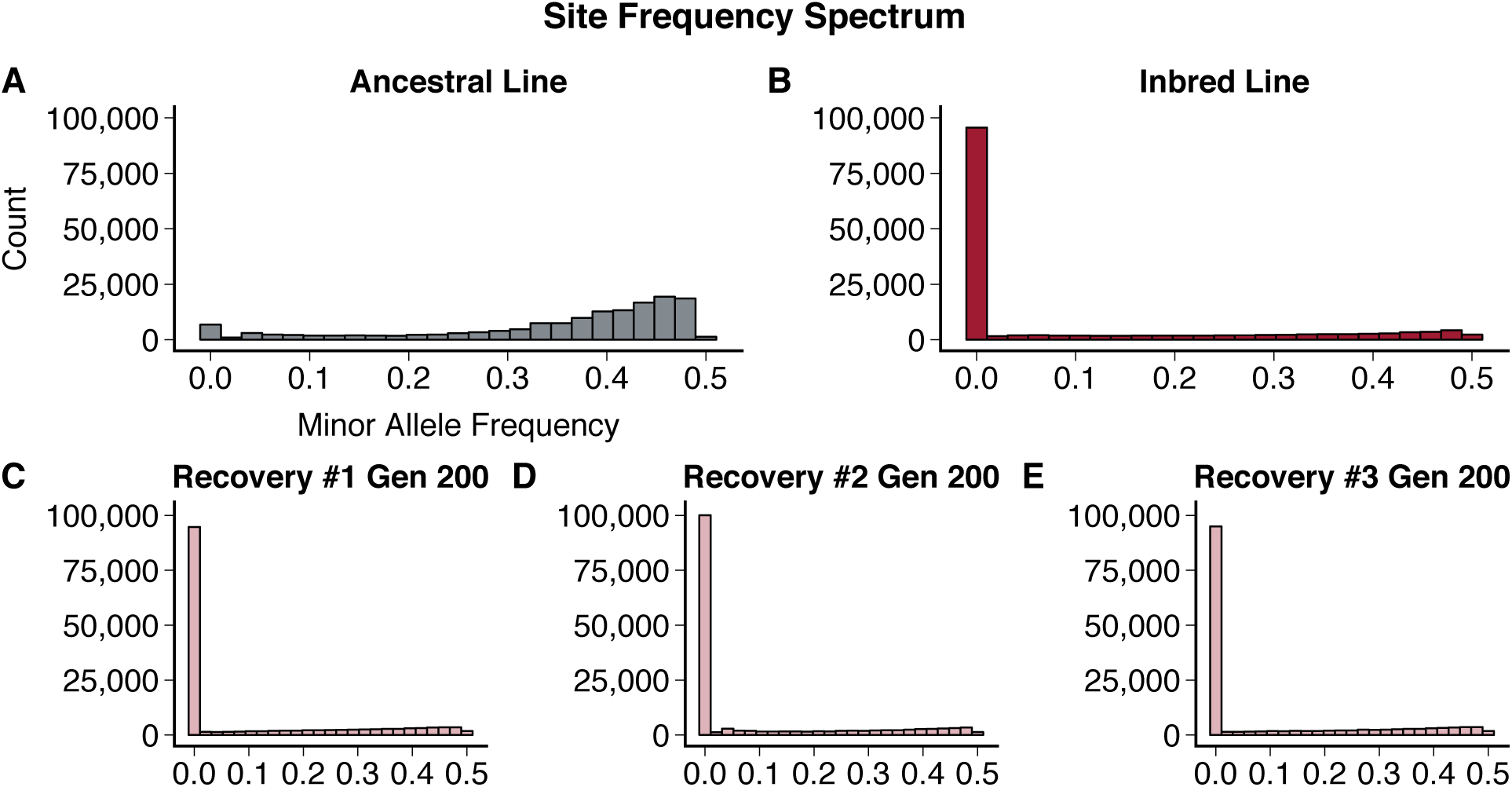
The minor allele site frequency spectrum showed (A) a majority of sites with minor allele frequencies 30-50% in the Ancestral line. This was altered through inbreeding and (B) the increase in fixation resulted in 98,940 fixed sites in the Inbred Line. Despite the intensity of inbreeding 48,490 sites still had segregating minor alleles. Recovery lines 1 (C), 2 (D), and 3 (E) had 9,394 shared sites retain fixation from the inbred line and 2,261 shared segregating minor alleles.

### Runs of Homozygosity

Heterozygosity peaks were larger in the Ancestor and smaller in the Inbred population but located in roughly similar regions (Fig. 4A-F). Chromosome X showed little change in heterozygosity after inbreeding (Fig. 5A, D) while Chromosome II showed a decrease in heterozygosity after inbreeding (Fig. 5B, E). Roughly one half of Chromosome IV showed a decrease in heterozygosity after inbreeding (Fig. 5C, F) while the second half retained heterozygosity through both inbreeding and recovery. The distribution of ROH increased in size and frequency in the Inbred line as compared with the Ancestor (SFig. 1).

**Figure 5.**
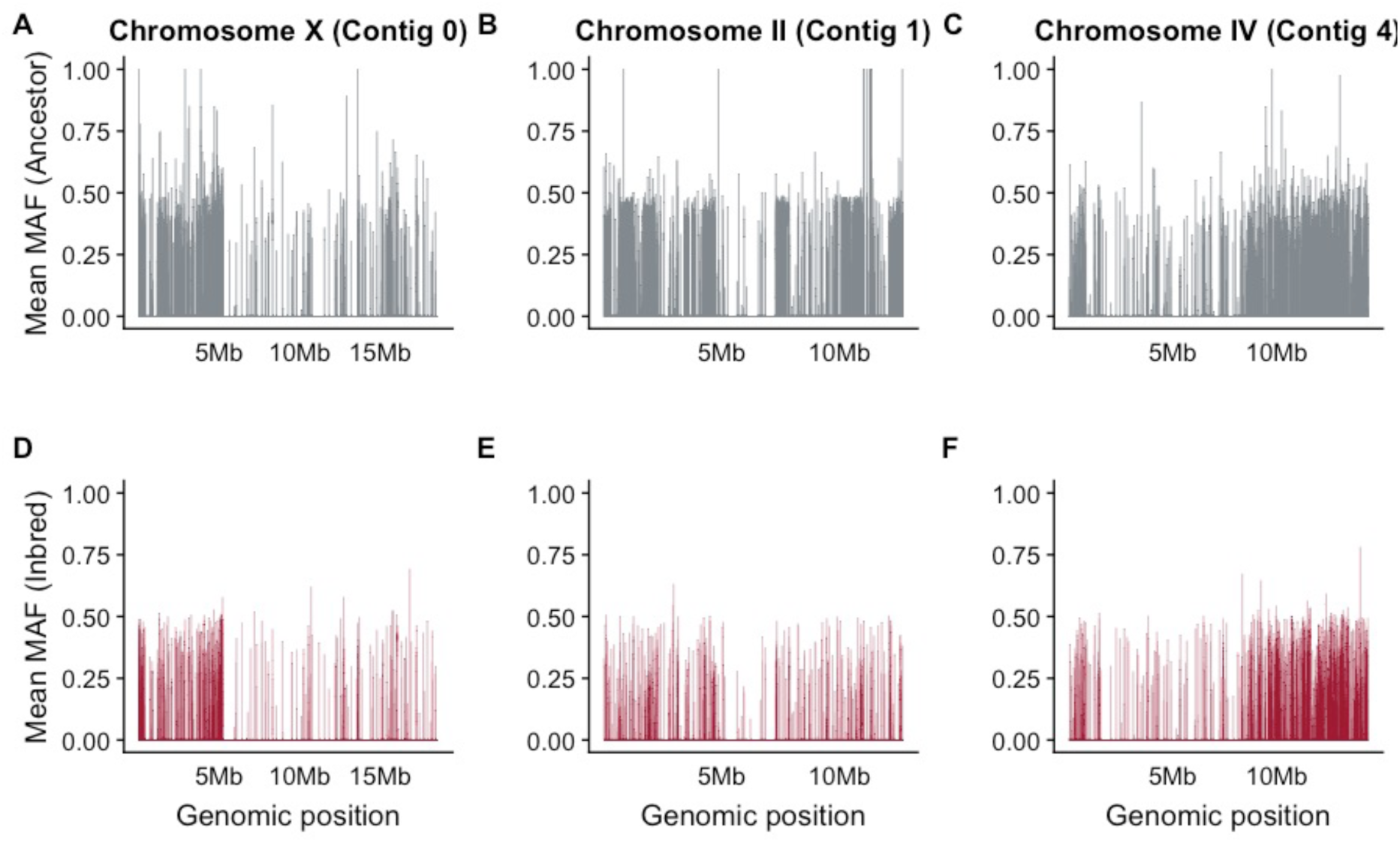
Runs of homozygosity across the 3 largest linkage groups (corresponding to (A) Chromosomes X, (B) II and (C) IV) show that polymorphism in the Ancestor line was decreased through inbreeding but regions of segregating variation remained in the Inbred line (D-F). Residual segregating polymorphisms are not evenly distributed along chromsomes and there are distinct regions of Chromosome X and IV that retain polymorphism in the Inbred line.

### Allele Frequency Trajectories

Of the 150,348 variable sites, 98,160 (65.29%) were segregating in the Ancestor and fixed during inbreeding. These sites were classified as ‘Fixation’ (Fig. 6A). Of these, 46,267 (30.77%) were at ‘Intermediate’ frequencies throughout inbreeding and recovery (Fig. 6B). The remaining 5,918 (3.94%) segregating sites were classified into trends based on their behavior during inbreeding and recovery. A small proportion of sites (3,868; 2.57% of the total variation) ‘bounced down,’ where the major allele frequency began high in the Ancestor, dropped during inbreeding, and increased during recovery (Fig. 6C). ‘Bounce up’ sites (343; 0.23% of the total) began at low frequency, rose during inbreeding, and decreased during recovery (Fig 5D). A small minority of sites (45; 0.035% of the total set) began at high frequency which was pushed down during inbreeding and continued to drop during recovery (Fig 5E). In 1,662 (1.11%) sites the major allele frequency rose during inbreeding and continued to rise during recovery (Fig 5F).

**Figure 6.**
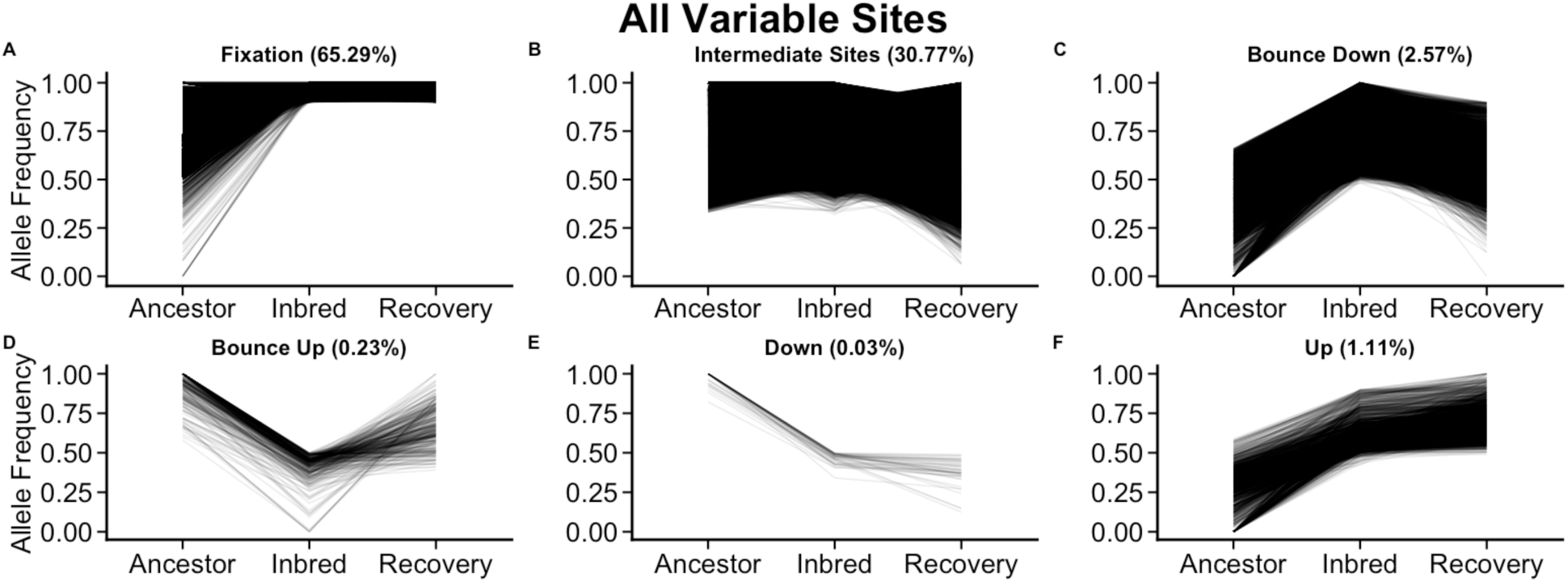
Across the entire genome allele frequency trajectories demonstrate that a majority of sites were either (A) fixed through inbreeding and remained fixed during recovery or (B) maintained intermediate allelic frequencies through both inbreeding and recovery. A minority of sites demonstrated allelic frequencies that were (C) low in the Ancestral line, raised through inbreeding and lowered again in the Recovery lines; (D) high in the Ancestral line, lowered through inbreeding and rose again in the Recovery lines; (E) lowered through inbreeding and lowered further in the Recovery lines; and (F) rose in frequency through inbreeding and rose further in the Recovery lines.

**Figure 7.**
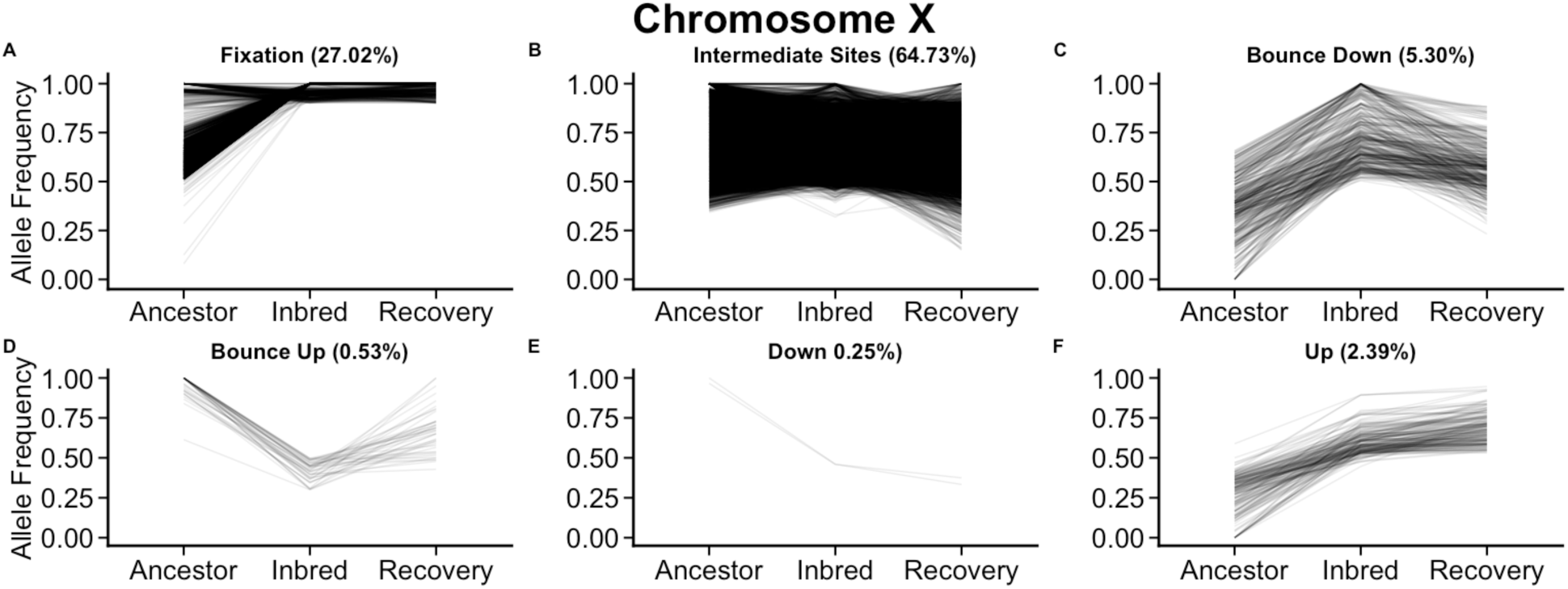
Nucleotides on the X Chromosome were less likely to (A) fix through inbreeding and (B) more likely to remain at intermediate frequency through inbreeding and recovery. A small proportion of sites on the X chromosome also showed parallel patterns of variable allele frequencies (C-F).

Nucleotides on the X chromosome had strikingly different patterns (Fig. 6G) with only 2,137 (27.02%) of the segregating nucleotides falling into our ‘Fixation’ scheme and 5,119 (64.73%) of sites segregating as ‘Intermediate.’ The remaining 670 (8.47%) sites showed Bounce Up, Down, Bounce Down, or Up patterns of segregation. In total 7,909 segregating sites (5.3% of the total set) resided on the X chromosome. The X chromosome is 18.6Mb and roughly 16% of the assembled 118.5Mb *C. remanei* genome. Segregating X sites were underrepresented in our analyses but displayed high genetic resistance.

### Effective Population Size

The effective population size of wild-collected *C. remanei* has been previously estimated to be ∼1,000,000 (Cutter et al. 2006). The poolSeq-estimated (Taus et al. 2017) effective population size was 26 for the Inbred line (S. Table 2). The three Recovery lines had a mean effective population size of 88 after 100 generations and 139 after 200 generations (S. Table 2).

### Selection Scans

The quasibinomial-GLM revealed 102 SNPs with significant parallel changes across the three Recovery lines (*q-value* < 0.05). Of these 102 SNPs, 36 were contained within 30 genes. Genomic locations and statistical estimation for these genes are given in Table 1. InterProScan protein domain annotations and *Caenorhabditis* orthologs for these genes are listed where available; several genes had no identifiable domain annotations or orthologous proteins in other species.

**Table 1.**
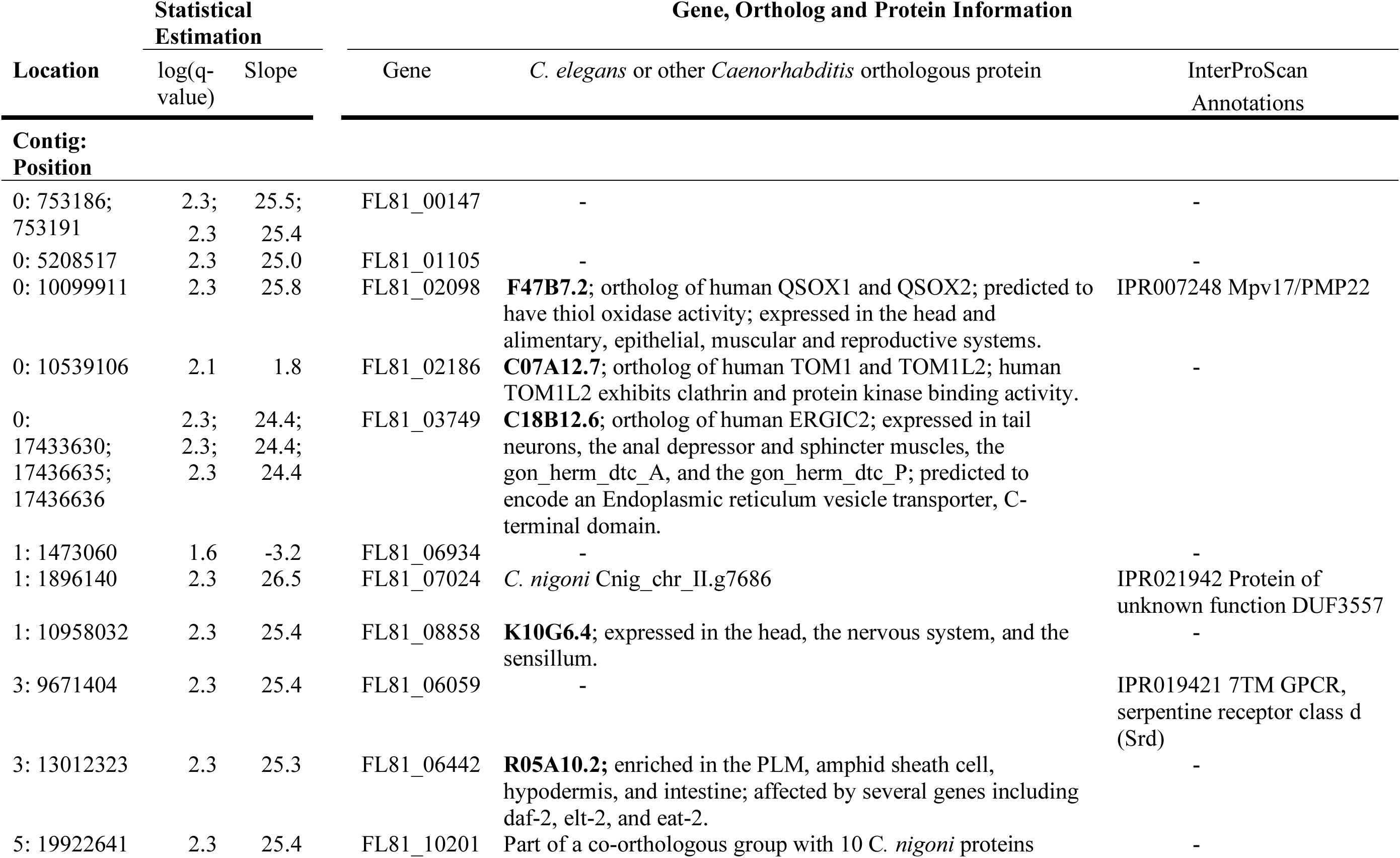
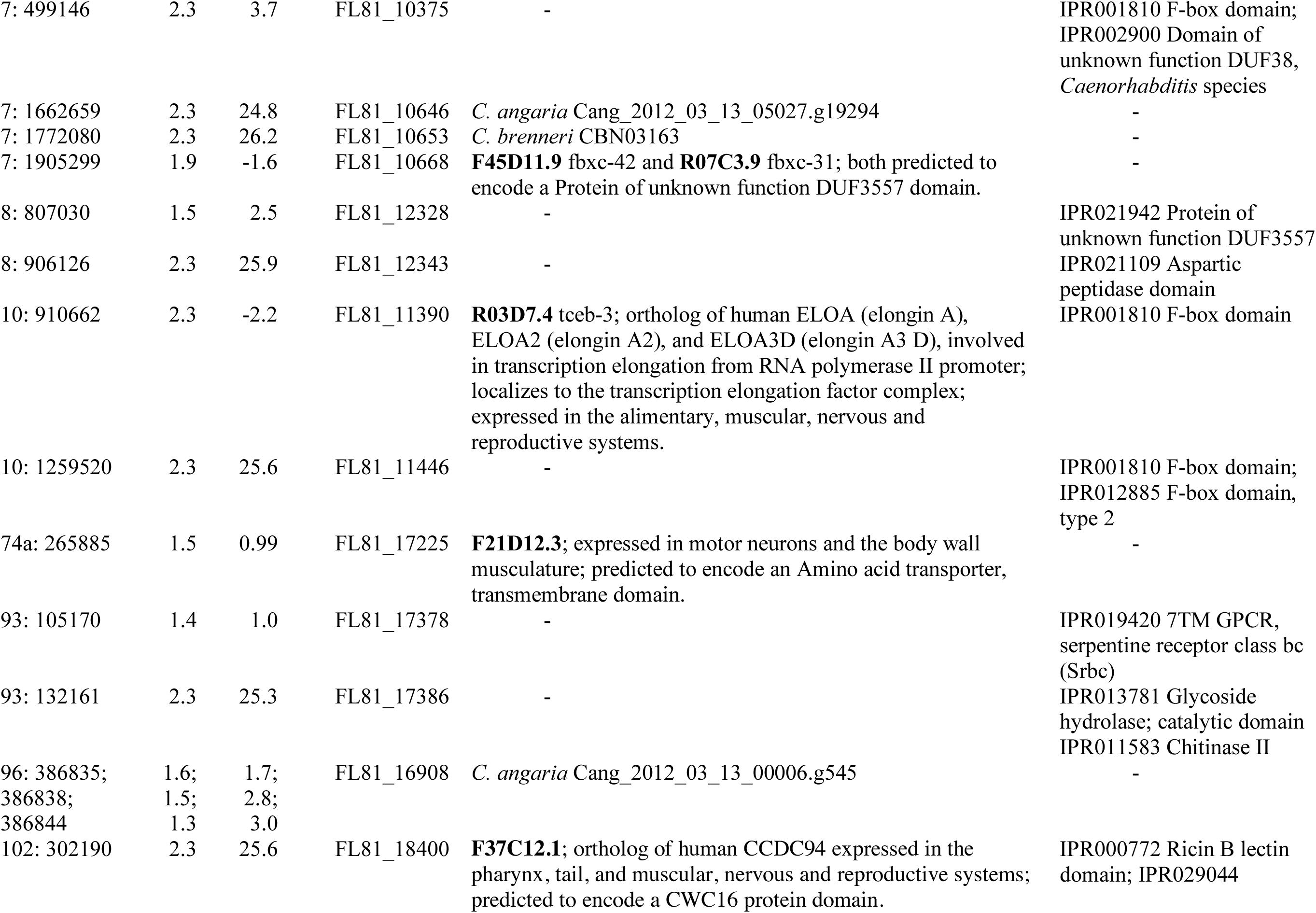
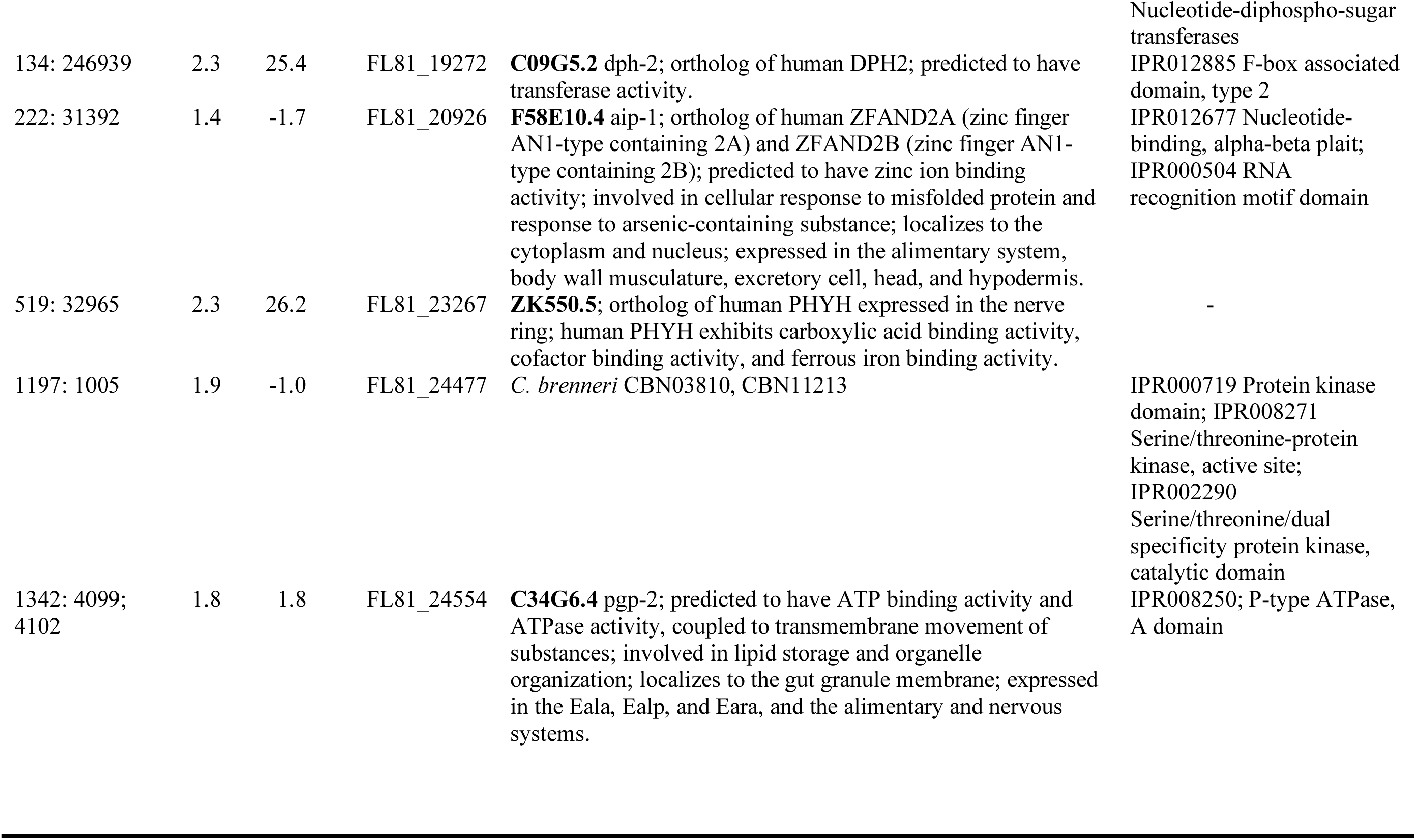
Genomic location, statistical estimation and gene name for each of the genes with significant SNPs in our allele frequency scans. Orthologous genes in *C. elegans* and other *Caenorhabditis* species and protein domain annotations are given where available.

### F_ST_

The mean per-site F_ST_ between Ancestor and Inbred lines was 0.5 and the distribution was strongly bimodal (Fig. 8). Roughly 30% of the variable sites in this comparison (74,505) had F_ST_ < 0.1 indicating little allelic divergence between the Ancestor and Inbred lines at these nucleotides. In contrast, ∼60% of the segregating sites had substantial F_ST_>0.5 between the Ancestor and Inbred.

**Figure 8.**
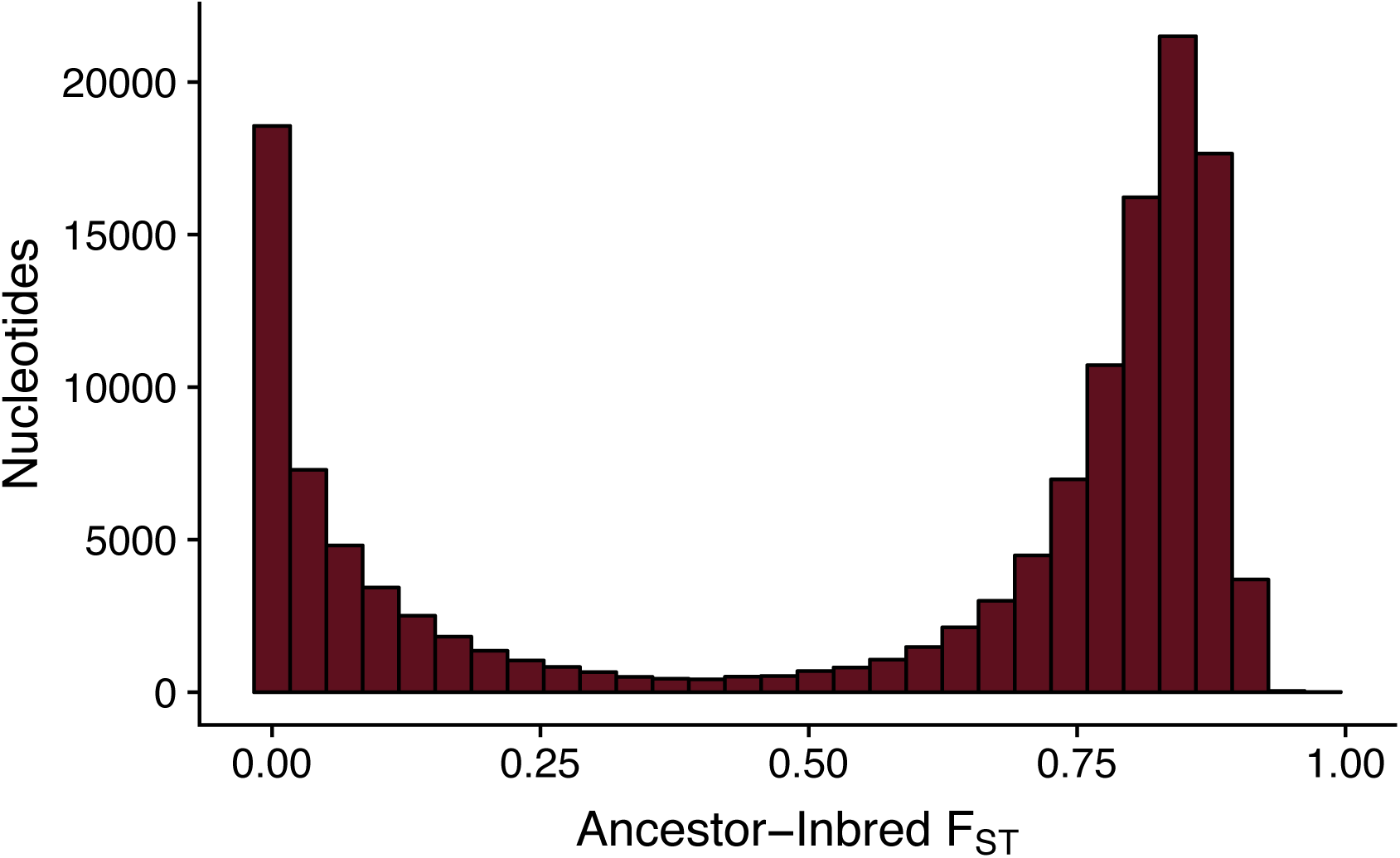
The frequency distribution of F_ST_ calculated between Ancestor and Inbred lines shows that there is a bimodal response to inbreeding with many nucleotides showing no divergence in allele frequency (i.e., F_ST_ ∼0) between Ancestor and Inbred lines and other sites showing high divergence in allele frequency in response to inbreeding (i.e., F_ST_ > 0.6).

## Discussion

The cycle of the generation of inbreeding depression and its subsequent recovery has probably been fundamentally important during the transition of breeding systems between outcrossing and self-fertilization (Charlesworth 2006), but at this moment in time is especially relevant to the future of species undergoing reductions in population size caused by human disturbance and global climate change (Gonzalez et al. 2013; Radchuk et al. 2019). While there is strong evidence from experimental populations that completely homozygous lines can indeed recover from fixed deleterious mutations (Burch and Chao 1999; Whitlock and Otto 1999; Estes and Lynch 2003), we find that the highly genetically diverse, outcrossing nematode *C. remanei* did not recover from inbreeding in our study. We found that 99% of strains died after just seven generations of inbreeding, and those that did survive had severely reduced fecundity (Fig. 4A). The fitness impacts of inbreeding are complemented by our genomic data, which show that the Inbred line had far fewer fixations than expected under a neutral model. Populations did not recover fecundity even after 300 generations of evolution at large population sizes. Overall, the severe reduction in fecundity with little recovery and complexity of the genomic response show that the effects of inbreeding are both detrimental and long lasting in *C. remanei*.

In contrast to our results, mutation accumulation studies have shown that it is possible to rapidly recover from complete homozygosity within experimental populations (Whitlock and Otto 1999; Maisnier-Patin et al. 2002; Estes and Lynch 2003; Burch and Chao 1999). Back-mutations at deleterious sites and beneficial mutations are thought to be rare (Smith 1978), but compensatory mutations may counteract fixed deleterious alleles and aid in fitness recovery (Whitlock and Otto 1999; Maisnier-Patin et al. 2002; Estes and Lynch 2003; Burch and Chao 1999). Mutation accumulation and recovery studies in *C. elegans* have demonstrated similar processes with compensatory epistatic mutations swept to fixation during recovery (Estes et al. 2011; Estes and Lynch 2003; Denver et al. 2010). For example, in a *C. elegans* mutation accumulation experiment 28 new mutations occurred during 60 generations of recovery (Denver et al. 2010). These mutations were subject to strong selective sweeps as they rose from undetectable to full fixation within 10-20 generations. Many of the new mutations had predicted interactions with well-characterized loci that had fixed during mutation accumulation, suggesting that these new mutations had compensatory beneficial effects.

Our results stand in stark contrast with these previous studies. There are several possible explanations for the difference in our results. First, it is possible that the landscape for compensatory mutations might differ across the species. While this seems extremely unlikely, it is a formal possibility that our data cannot directly address. More likely is a difference in how compensatory mutations interact with differences in mating systems between *C. elegans* and *C. remanei.* Under self-fertilization in *C. elegans*, compensatory mutations that arise in a given genetic background, even if they are on a different chromosome, are very likely to be inherited with the target deleterious mutation because, although recombination does occur, it has little effect on genetic diversity when the rest of the genome is nearly completely homozygous. In contrast, obligate outcrossing in *C. remanei* increases the effectiveness of recombination in breaking up different genetic combinations, especially in large populations. This may make it more difficult for epistatically interacting loci to remain together on the same genetic background (Phillips 2008). On the other hand, in *C. elegans* other deleterious mutations that are not “fixed” by the compensatory mutation are locked in the genome, whereas in *C. remanei*, different combinations of adaptive mutations can recombine into a common background much more easily, which should be relevant on the timescales of this study. More importantly, since our experiments were initiated from a highly inbred state, recombination would have little impact on changing the dynamics of deleterious mutations that are already fixed in the population, since they would be present on every genetic background upon which a new compensatory mutation might find itself. Overall, then, while differences in genetic systems in species used in mutation accumulation and our genetic recovery experiments could explain some of the differences in results, they are unlikely to explain the extreme difference in rate of total fitness recovery across approaches.

The most likely cause of the differences observed here are differences in the genetic architecture of segregating mutations under inbreeding depression and novel mutations under mutation accumulation. There are three main difference here. First, because mutation accumulation experiments are designed to capture as many mutations as possible by reducing the effective population size of each experimental line to be as small possible (*N* = 1 in the case of *C. elegans*), the main effects of mutations in mutation accumulation experiments might be much larger than those that escape natural selection within segregating populations. Similarly, we would expect most of the variants fixed during the generation of inbreeding depression to be recessive (Charlesworth and Charlesworth 1999), whereas mutations in mutation accumulation studies can in principle have any dominance effects (albeit with some bias toward recessivity). These two factors make it much more likely that the main mutational effects “fixed” by compensatory mutations in a mutation-accumulation recovery experiment will have larger effects than most segregating variation under inbreeding depression, which might make them more likely targets for compensatory change.

However, the third and most likely explanation based on genetic architecture for the extremely slow recovery of fitness under inbreeding—and the one most clearly supported by the genomic data—is that there are simply many more segregating deleterious mutations in natural populations than are generated in mutation-accumulation experiments. Our “ancestral” *C. remanei* population displayed high levels of polymorphism at many different sites. In particular, generation of the initial inbred line revealed the presence of many recessive lethal alleles under close interbreeding (Fierst et al. 2015; see also Dolgin et al. 2007). The surviving inbred population had a severe loss of fitness due to fixation of many slightly deleterious alleles. The presence of numerous sites that are actually resistant to complete inbreeding suggests that *C. remanei* populations are subject to high levels of segregation load and carry complex incompatible genetic combinations. The complex structure of the genetic load of the ancestral *C. remanei* population was therefore likely critical to the constrained recovery demonstrated in our Recovery lines.

Despite the constrained recovery in fitness, there was clearly very strong and consistent selection for alleles leading to evolutionary rescue via new mutations. We were still able to detect 102 SNPs with parallel changes across the three Recovery lines. 36 of these sites were found within 30 genes, and we were able to determine some functional information for many of these genes (Table 1). The majority are involved in alimentary, muscular, nervous and reproductive systems. Given the low fitness recovery we observed and the complexity of gene interactions (Phillips 2008) these parallel changes indicate alleles with strong phenotypic effects. So, we do in fact see a clear signal for an evolutionary response, but it is spread across many different independent sites. Many, many more sites display independent response within in each replicate, and many of these are likely to be functional relevant, however it is difficult to distinguish these from other possible effects, including genetic drift, without more formal functional validation. These genes, and the alleles we identified in the Recovery lines, are potential targets for molecular manipulation and CRISPR genome editing for studying genotype-phenotype-fitness relationships in *C. remanei*.

### Deleterious mutations and aging

Unlike fecundity, lifespan did not show any decrease under inbreeding. Instead, the Recovery lines evolved an increase in lifespan when compared with both the Ancestor and Inbred lines (Fig. 2B). The basic premise of inbreeding depression is traits decline in value because deleterious alleles will always have a negative effect on traits under positive directional selection. A lack of decline in longevity with inbreeding would therefore suggest that longevity itself is not under selection, nor is it strongly correlated with other traits under selection. This result is consistent with an experimental evolution study in *C. elegans* which did not find any evidence for a tradeoff between early reproduction and longevity (Anderson et al. 2011). Alternatively, the alleles involved in lifespan extension could have been physically or statistically linked to a region under selection in the Recovery lines. We did identify parallel allelic changes in FL81_06442, a *C. remanei* protein orthologous to the *C. elegans* protein R05A10.2. This protein is affected by *daf-2*, an aging factor, in *C. elegans* (Kenyon et al. 1993) and may be a target for further studies investigating lifespan in *C. remanei*.

### Genetic basis of inbreeding depression

Our genomic data showed that fixation and resistance to inbreeding were not consistent across the genome. The X chromosome in particular showed genetic resistance with 73% of variable sites retaining ancestral polymorphism after inbreeding. In *C. remanei*, as in other Rhabditid nematode species, females carry 2 X chromosomes (denoted XX) and males carry a single X chromosome (denoted X0) with no Y or male-specific chromosome (Brenner 1974; Nigon and Dougherty 1949). This exposes the X chromosome to different selection dynamics since recessive deleterious alleles are exposed in haploid condition in males and may have already been purged by purifying selection prior to inbreeding. High levels of genetic resistance on the X chromosome may also imply that *C. remanei* genetic load and resistance to inbreeding are related to sex-specific selection and X-autosome epistasis that differs for males and females.

Sexually reproducing organisms are expected to accumulate extensive suites of mildly deleterious loci when found at large population sizes that can lead to substantial inbreeding depression when shifted to smaller population sizes. In a sense, the change in fitness due to inbreeding is not qualitatively different to changes in the environment in which previously favorable alleles are now deleterious. In both cases, new mutations are needed to allow the species to escape the new state of low fitness in order to adapt and escape the possibility of eventual extinction. Given the rapidly changing face of the planet, there are has been recent renewed attention to the importance of “evolutionary rescue” as a means of confronting continuing degradation of the environmental and genetic landscape (Bell 2019). Despite some hopeful indications based on earlier mutation-accumulation studies, our results indicate that evolutionary rescue alone may not be powerful enough for recovery from inbreeding (Stewart et al. 2017). For the nematode *C. remanei*, this is almost certainly caused by the very large number of segregating deleterious alleles in the population prior to inbreeding. The total number of loci involved makes it impossible for a small number of compensatory mutations to lead to rapid recovery of fitness. Part of the complexity of the genetic basis of inbreeding depression in this species is due to the very large effective population sizes at which it exists in nature. It is possible that species with smaller population sizes might have few segregating alleles before inbreeding, leading to less severe fitness effects. On the other hand, those species are also likely to exist at large enough population sizes to allow a sufficient number of compensatory mutations to enter the population before demographic factors drive the population to extinction. Overall, our results suggest that evolution is unlikely to lead to rapid rescue of endangered populations, at least from a genetic point of view.

## Acknowledgments

We gratefully acknowledge the helpful feedback and comments from members of the Fierst lab. This research was conducted with Government support under and awarded by DoD, Air Force Office of Scientific Research, National Defense Science and Engineering Graduate (NDSEG) Fellowship, 32 CFR 168a to PEA and NIGMS GM102511 to PCP.

## Author Contributions

PCP conceived the experimental study, ALC and EMY conducted the experimental study, and JHW conducted the genomic sequencing. JLF and PEA conceived and conducted the analyses. PEA, JLF and PCP wrote the initial manuscript and all authors contributed to and reviewed the final manuscript.

## Data Accessibility

Whole genome sequence data associated with this project have been deposited with the National Center for Biotechnology Information under BioProject PRJNA562722. All bioinformatic scripts and workflows are accessible at https://github.com/Bamacomputationalbiology/Inbreeding.

